# Structural similarities between some common fluorophores used in biology and marketed drugs, endogenous metabolites, and natural products

**DOI:** 10.1101/834325

**Authors:** Steve O’Hagan, Douglas B. Kell

## Abstract

**Background:** It is known that at least some fluorophores can act as ‘surrogate’ substrates for solute carriers (SLCs) involved in pharmaceutical drug uptake, and this promiscuity is taken to reflect at least a certain structural similarity. As part of a comprehensive study seeking the ‘natural’ substrates of ‘orphan’ transporters that also serve to take up pharmaceutical drugs into cells, we have noted that many drugs bear structural similarities to natural products. A cursory inspection of common fluorophores indicates that they too are surprisingly ‘drug-like’, and they also enter at least some cells. Some are also known to be substrates of efflux transporters. Consequently, we sought to assess the structural similarity of common fluorophores to marketed drugs, endogenous mammalian metabolites, and natural products. We used a set of some 150 fluorophores.

**Results:** The great majority of fluorophores tested exhibited significant similarity (Tanimoto similarity > 0.75) to at least one drug as judged via descriptor properties (especially their aromaticity, for identifiable reasons that we explain), by molecular fingerprints, by visual inspection, and via the “quantitative estimate of drug likeness” technique. It is concluded that this set of fluorophores does overlap a significant part of both drug space and natural products space. Consequently, fluorophores do indeed offer a much wider opportunity than had possibly been realised to be used as surrogate uptake molecules in the competitive or trans-stimulation assay of membrane transporter activities.

## INTRODUCTION

Fluorescence methods have been used in biological research for decades, and their utility remains unabated (e.g. [1–16]). Our specific interest here is in the transporter-mediated means by which small fluorescent molecules enter living cells, and our interest has been stimulated by the recognition that a given probe may be a substrate for a large variety of both influx and efflux transporters [17]. Efflux transporters are often fairly promiscuous, since their job is largely to rid cells of unwanted molecules that may have entered, although they can and do have other, important physiological roles (e.g. [18–28]), and they are capable of effluxing a variety of fluorescent probes (e.g. [29–36]). However, given that most of these probes are contemporary, synthetic molecules, the uptake transporters for which they are substrates must have evolved in nature for other purposes. These purposes may reasonably be expected to include the uptake of endogenous metabolites in multicellular organisms [37–40], as well as exogenous natural products whose uptake can enhance biological fitness (e.g. [41; 42]). This explanation does seems to hold well for synthetic, marketed pharmaceutical drugs [42].

Consequently, it seemed reasonable that existing fluorescent molecules, not specifically designed for the purpose but that are taken up by biological cells, might also bear structural similarities to endogenous substrates (metabolites) and to natural products, and potentially also to marketed drugs. If so, they might then serve as surrogate transporter substrates for them. Indeed, there are examples - so-called fluorescent false neurotransmitters - where such fluorescent analogues of natural substrates have been designed precisely for this purpose (e.g. [43; 44]). The aim of the present work was to assess the extent to which this kind of structural similarity between (i) common fluorophores used in biology and (ii) other molecular classes (endogenous mammalian metabolites, marketed pharmaceutical drugs, and known natural products) might be true. It is concluded that in structural terms common fluorophores do indeed overlap drug space significantly, and we offer an explanation based on the consonance between aromaticity, conjugated π-bonds, and fluorescence.

## MATERIALS AND METHODS

Fluorophores were selected from the literature and by scanning various catalogues of fluorophores, and included well known cytochemical stains, food dyes, laser dyes and other fluorophores, including just a few marketed drugs plus fluorescent natural products. We chose only those whose structures were known publicly. The final set included 150 molecules. Supplementary Table 1 gives a spreadsheet of all the relevant data that we now discuss, including the marketed drugs, Recon2 metabolites [45] (both given also in ref 37) and a subset of 2000 natural products from UNPD (see [42; 46]).

Although there are a great many possible molecular encodings (whether using molecular fingerprints or vectors of calculated properties), each of which can give a different Tanimoto similarity, for our present purpose we chose to use only the Patterned encoding within RDKit (www.rdkit.org/). We also used the RDKit version of QED (https://www.rdkit.org/docs/source/rdkit.Chem.QED.html). Workflows were written in KNIME as per our standard methods [37-40; 42; 47-49]. t-SNE plots used the first 10 PCs (95.3% explained variance) as inputs based on 27 RDKit descriptors, and were otherwise as previously described [50].

## RESULTS

Figure 1A gives a Principal Components Analysis (PCA) plot of the distribution of the four classes based on a series of descriptors in RDKit (www.rdkit.org/), while Fig 1B gives a t-SNE [51] plot of the same data. These clearly show a strong overlap between the rather limited set of fluorophores used and quite significant parts of drug space. Fig 1C gives just the fluorophores, with the nominal excitation maximum encoded in its colour. This suggests that even with just ~150 molecules we have achieved a reasonable coverage of the relevant ‘fluorophore space’, with no obvious bias, nor trend in excitation wavelengths.

**Figure 1.**
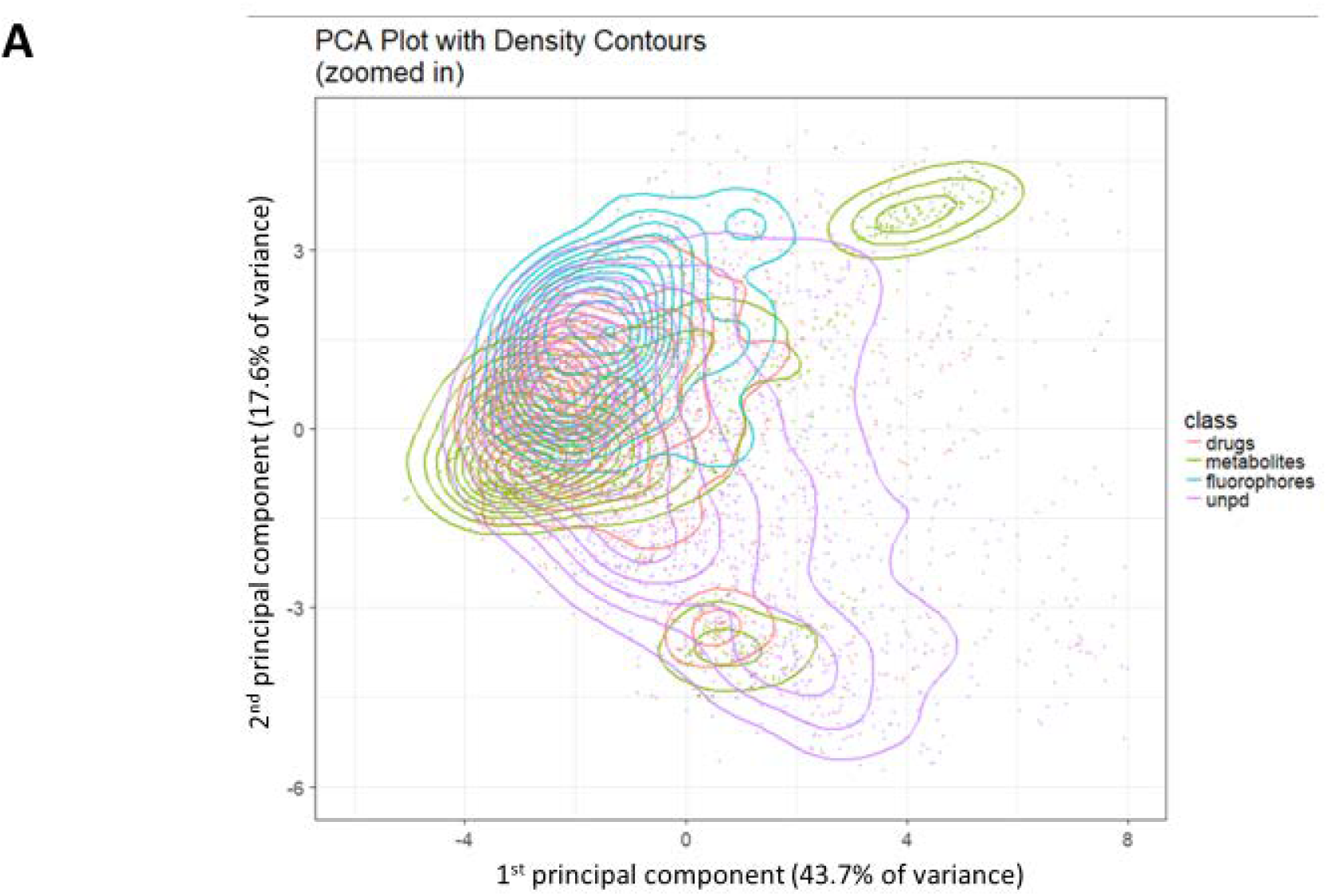
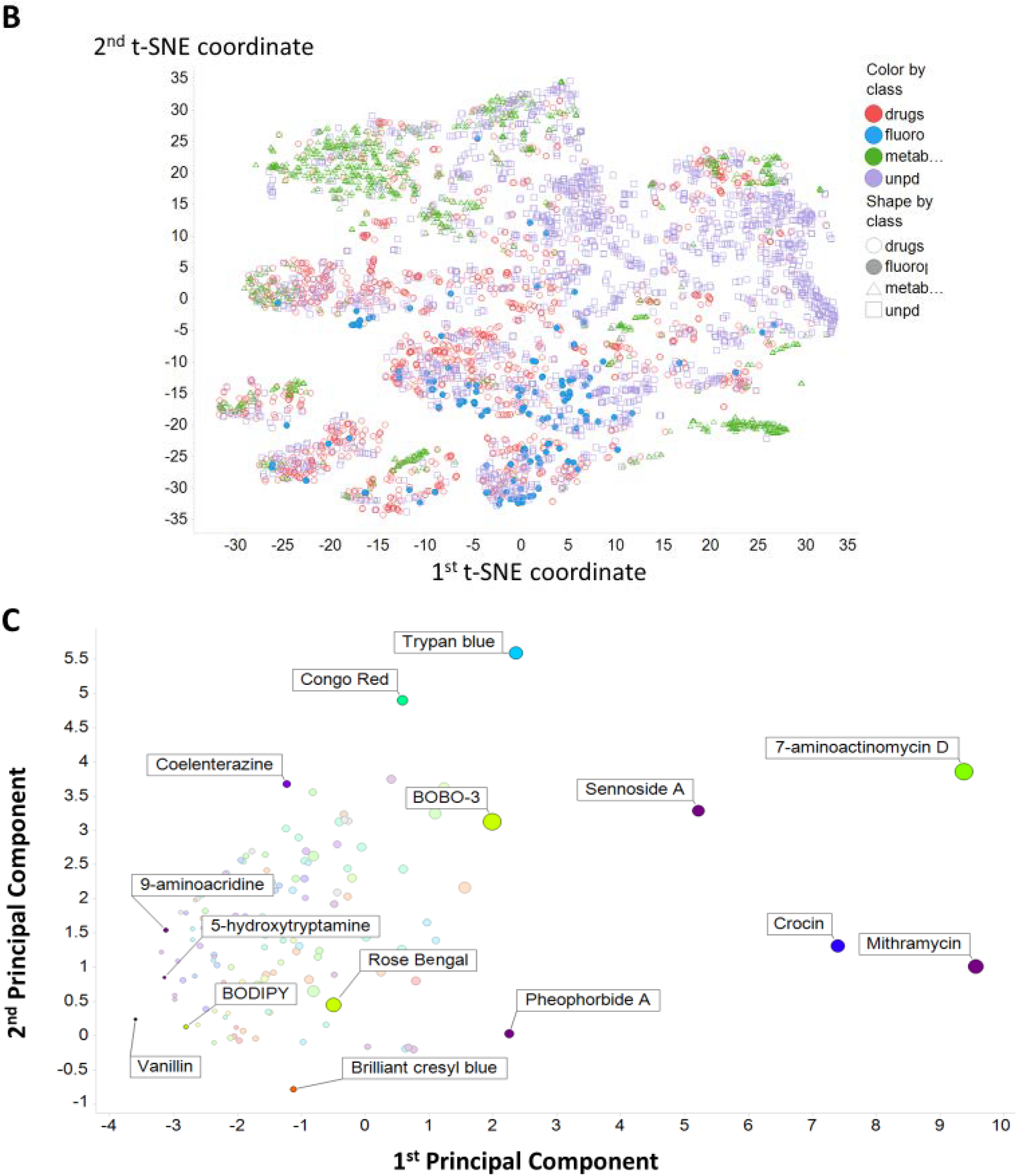
Principal components and t-SNE plots of the principal components of the variance in calculated properties of the molecules used. A. The first two principal components of the variance in calculated properties of the four classes fluorophores, drugs, metabolites and natural products. Molecules are as in Table S1, with drugs and metabolites being those given in [37]. A sampling of 2000 natural products from our download [42] of UNPD was used. Descriptors were z-scores normalised and correlation filtered (threshold 0.98. B. t-SNE plot of the data in (A), using the same colour coding. C. Plot of the first two principal components of the variance of the fluorophores alone. The excitation wavelength is encoded in the colour of the markers. The size of the symbol encodes the molecular weight, indicating that much of the first PC is due to this (plus any other covarying properties).

We previously developed the use of rank order plots for summarising the relationships (in terms of Tanimoto similarities) between a candidate molecule or set of molecules and a set of targets in a library [37]. Fig 2 shows such a rank order plot, ranking for each fluorophore the most similar molecule in the set of endogenous Recon2 [37; 52] metabolites, the set of marketed drugs [37], and a random subset of 2000 of some 150,000 molecules taken [42; 49] from the Unified Natural Products Database (UNPD) [46]. This again shows very clearly that the majority of fluorophores chosen do look moderately similar (TS>0.75) to at least one drug (and even more so to representatives of the natural products database).

**Figure 2.**
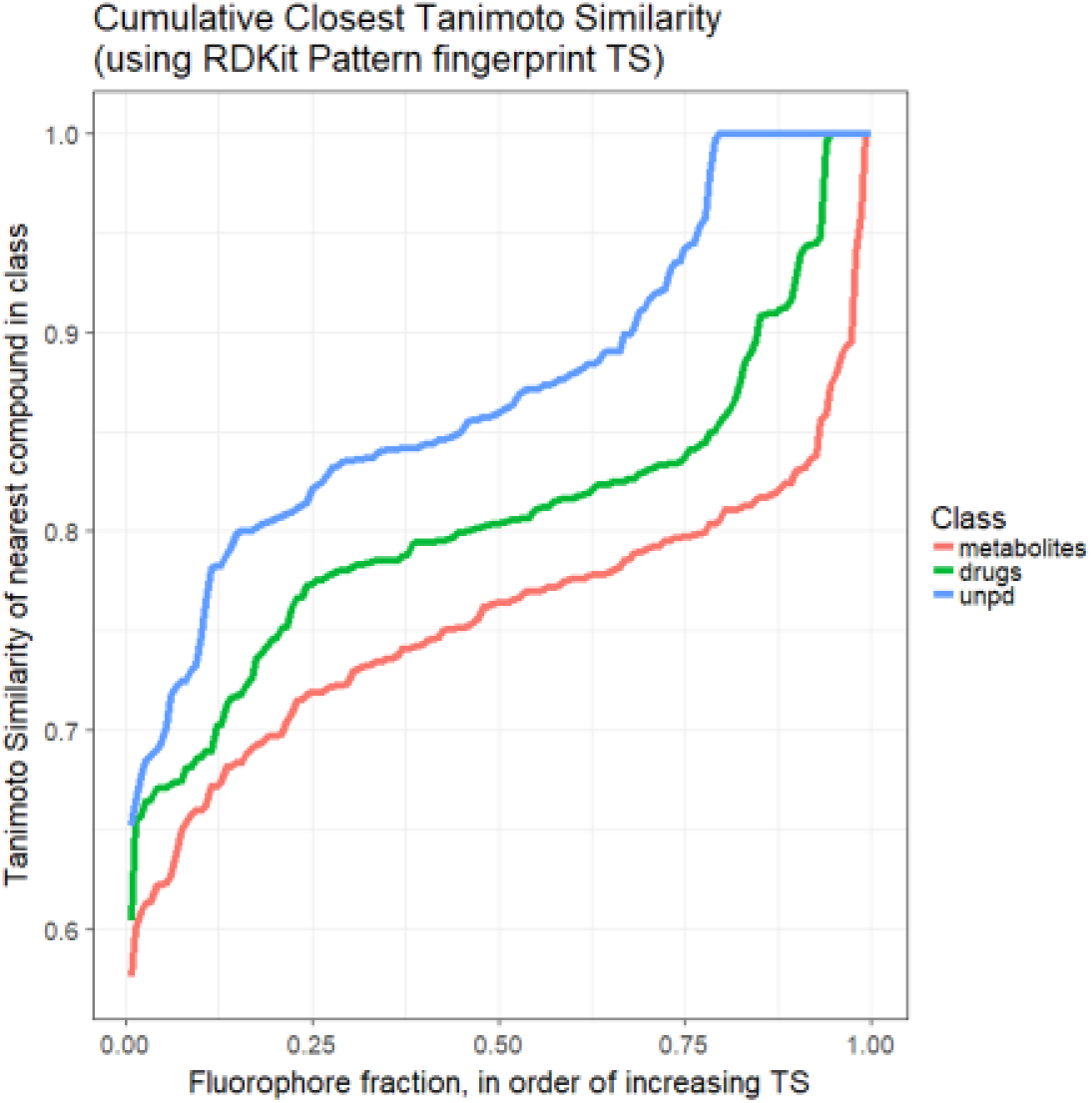
Ranked order of Tanimoto similarity for fluorophores vs marketed drugs 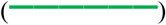, fluorophores vs Recon2 metabolites 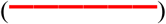, and fluorophores vs a 2000-member sampling of UNPD 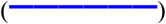. Each fluorophore was encoded using the RDKit ‘Patterned’ encoding, then the Tanimoto similarity for it calculated against each drug, metabolite or natural product sample. The highest value of TS for each fluorophore was recorded and those values ranked. Read from right to left.

It is also convenient [37] to display such data as a heat map [53], where a bicluster is used to cluster similar structures and the colour of the cell at the intersection encodes their Tanimoto similarity. Figure 3 shows such heatmaps for (A) fluorophores vs endogenous (Recon2 [45]) metabolites, (B) drugs, and (C) 2000 sampled natural products from UNPD. The data reflect those of Fig 2, and it is again clear that for each fluorophore there is almost always a drug or a natural product for which the average Tanimoto similarity is significantly greater than 0.7.

**Figure 3.**
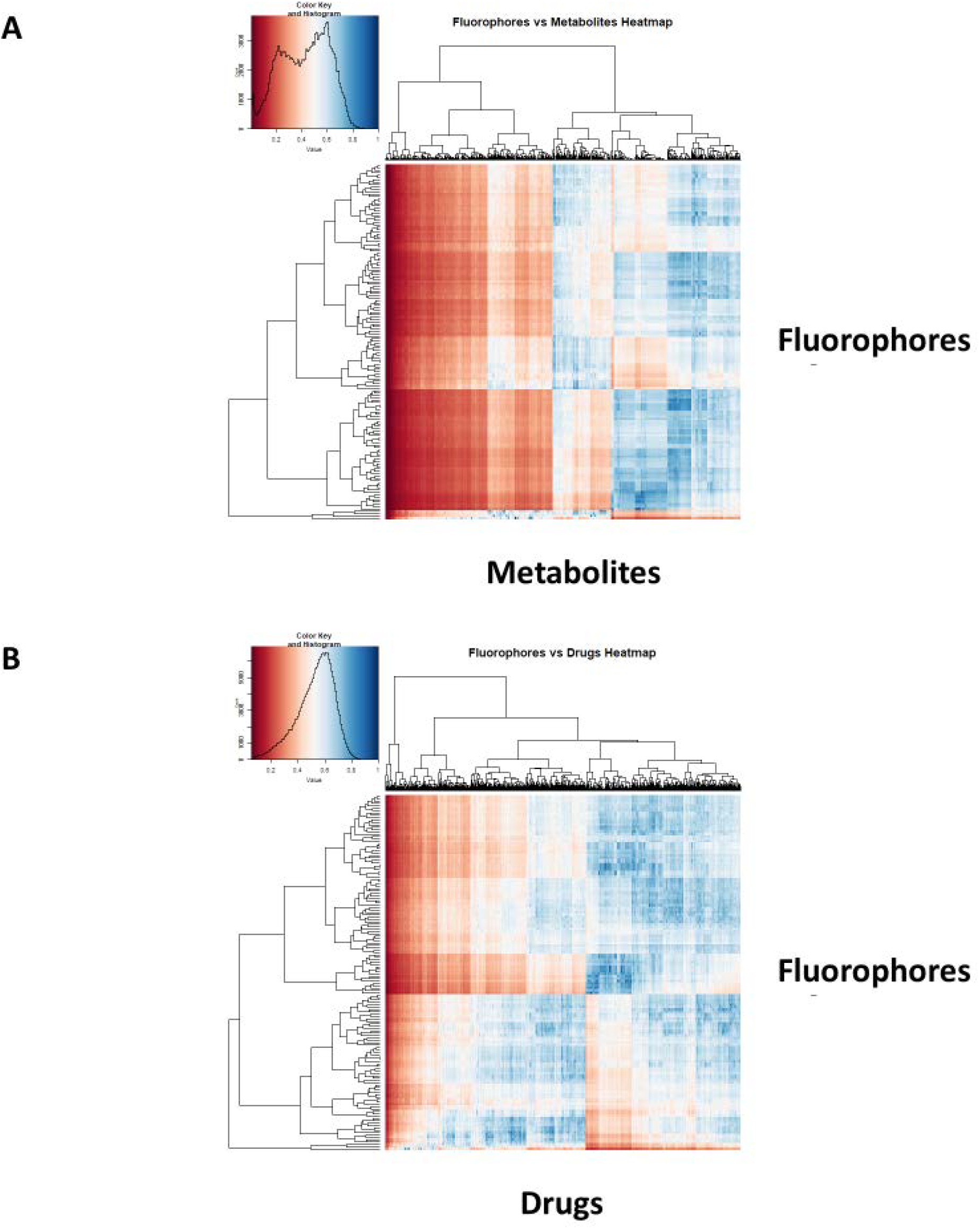
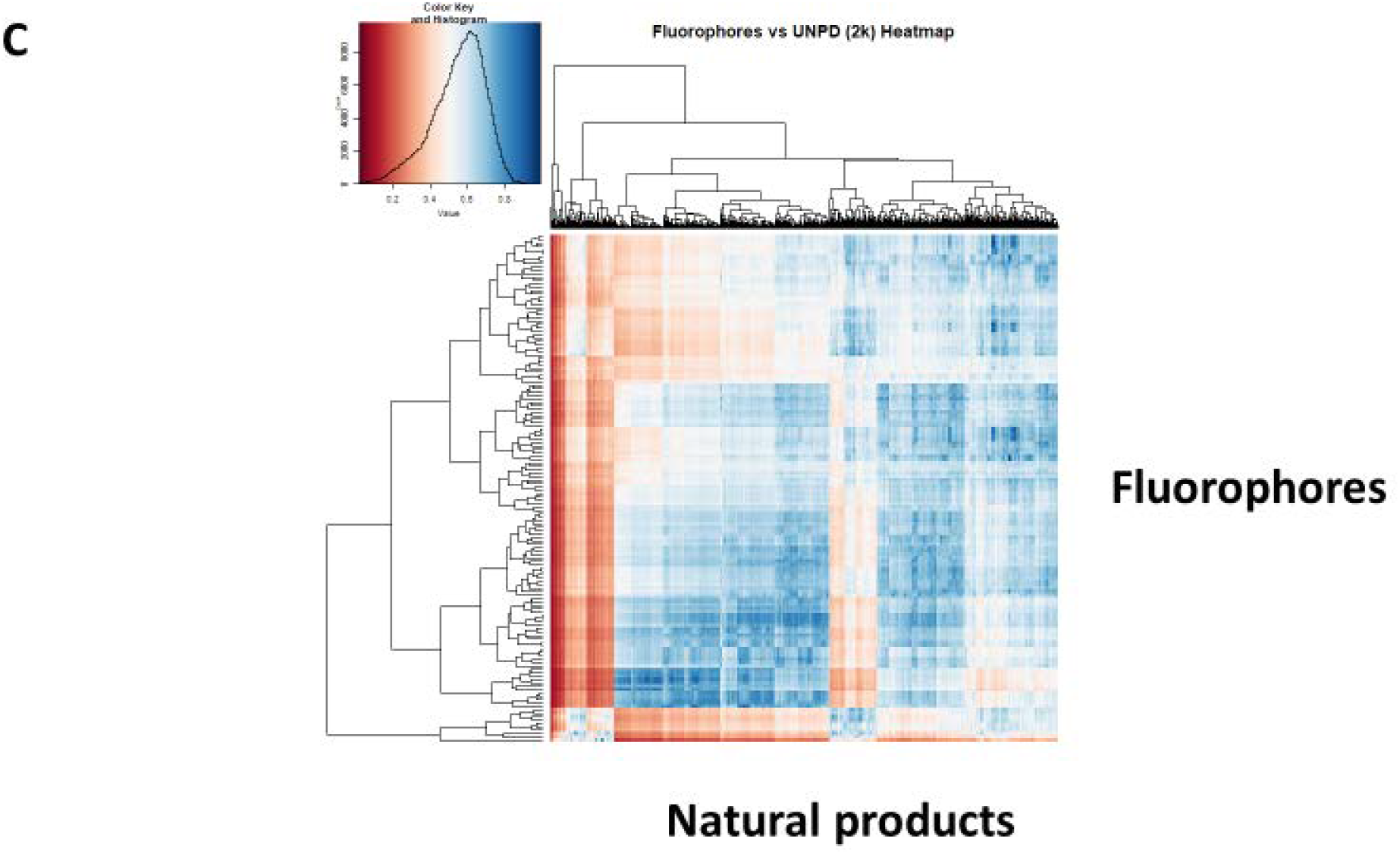
Heat maps illustrating the Tanimoto similarities (using the RDKit patterned encoding) between our selected fluorophores and (A) Recon2 metabolites, (B) Drugs, and (C) a subset of 2000 natural products from UNPD.

While is rather arbitrary, to say the least (given how the Tanimoto similarity varies with the encoding used), as to whether a particular chemical structure is seen by humans as ‘similar’ to another, we provide some illustrations that give a feeling for the kinds of similarity that may be observed.

Thus (Fig 4A) we illustrate the drugs closest to fluorescein in t-SNE space (as per Figure 1B), since fluorescein is a very common fluorophore, is also widely used in ophthalmology (e.g. [54; 55]), and can enter cells via a variety of transporters [56] such as monocarboxylate transporters (SLC16A1, SLC16A4) [57], SLC01B1/3B1 [58; 59] and SLC22A20 [60] (see also Table 1).

**Figure 4.**
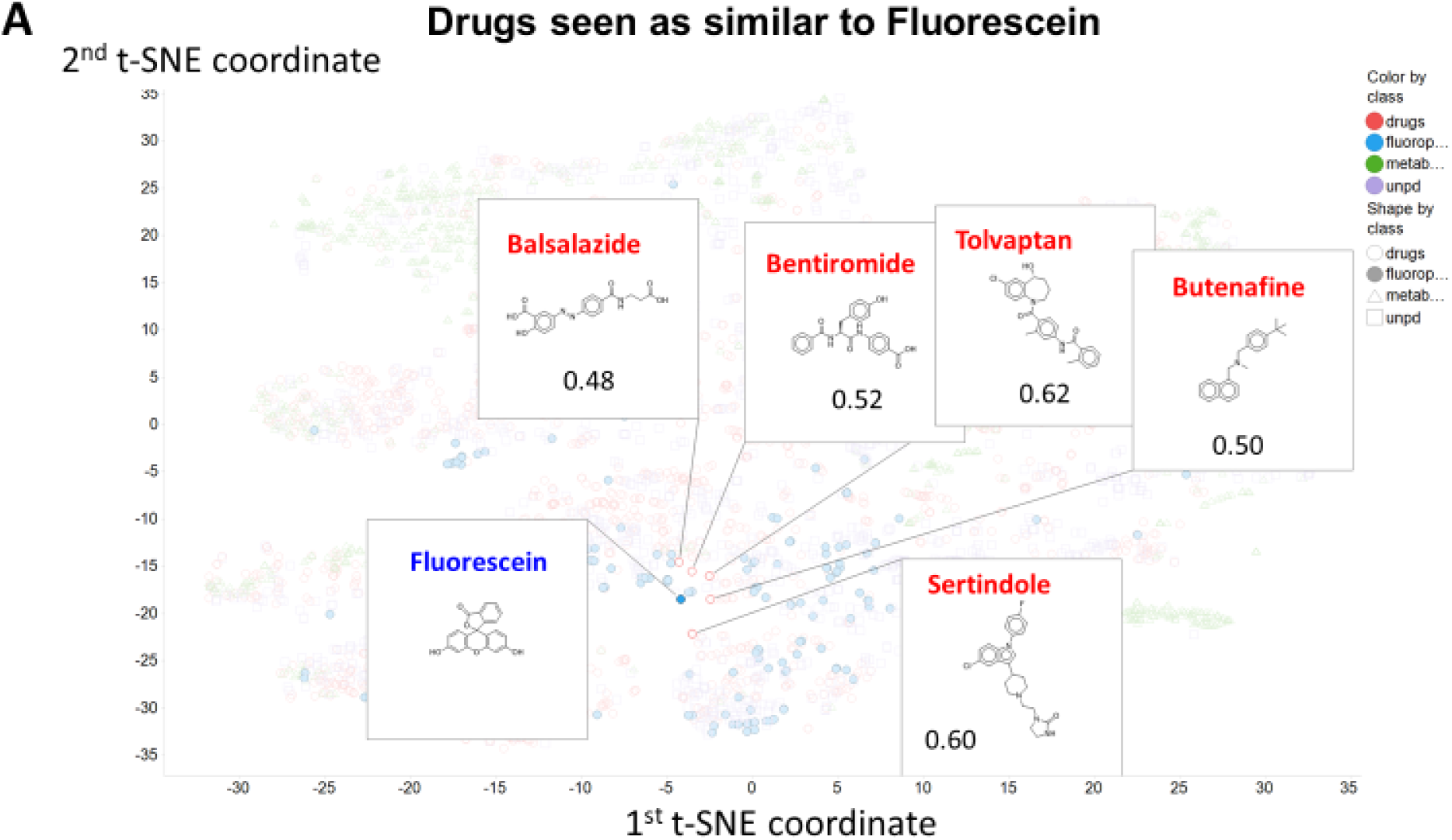
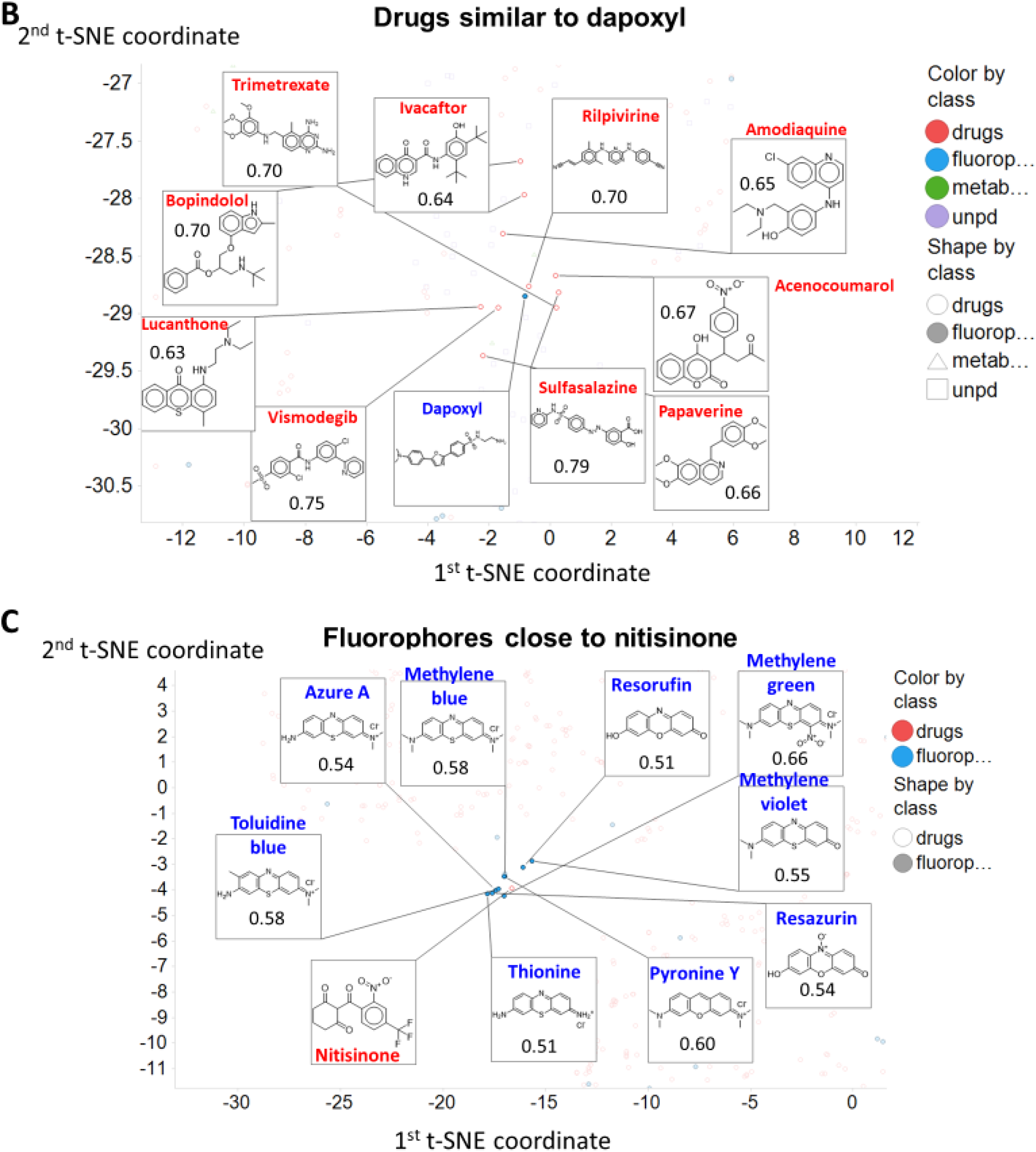
Observable structural similarities between selected fluorophores and drugs. The chosen molecules are (A) fluorescein, (B) dapoxyl (both fluorophores) and (C) nitisinone (a drug). Data are annotated and/or zoomed from those in Fig 1B.

**Table 1.** Some examples in which fluorescent dyes have been found to interact with uptake transporters directly as substrates or inhibitors. We do not include known non-fluorescent substrates to which a fluorescent tag has been added (see e.g. [100–102]).

| Dye | Transporter | Comments | Reference |
| --- | --- | --- | --- |
| Amiloride | OCT2 (SLC22A2) | Also a drug. Rhodamine 123 and 6G also served as substrates. | [103] |
| 4',6-diamidino-2-phenylindol (DAPI) | OCT1 (SLC22A1) | Potently inhibited by desipramine. Also by various organophosphate pesticides. | [104]; [105] |
| DiBAC(4)3 | Na<sup>+</sup>/HCO<sub>3</sub><sup>-</sup> NBCe1-A<br>SLC4A4 | Competes with 4,4'-Diisothiocyanatostilbene-2,2'-disulfonic acid | [106] |
| 5-carboxyfluorescein | OAT3 (SLC22A8) | Very high V<sub>max</sub> | [60] |
| 6-carboxyfluorescein | OAT1 (SLC22A6) |  | [60]; [107] |
| 2',7'-dichlorofluorescein | OATP1B1 (SLCO1B1) | Good substrate | [108] |
| 4-(4-(Dimethylamino) styryl)-N-methylpyridinium (ASP<sup>+</sup>) | Dopamine transporter (SLC6A3) |  | [109; 110] |
|  | Noradrenaline transporter (SLC6A2) |  | [109; 111; 112] |
|  | Serotonin transporter SLC6A4 |  | [109; 113] |
|  | Various monoamine transporters |  | [114] |
|  | OCT1/OCT2 (SLC22A1/2); | Seen as a model substrate | [115] |
|  | Various OCT transporters |  | [116] |
|  | Other, unknown (non-OCT1/2) transporters with low affinity |  | [117] |
| Ethidium | OCT1/2/3 (SLC22A1/2/3) | Substrate | [118] |
| FFN511 | VMAT2 (SLC18A2) | 'False fluorescent neurotransmitter' (i.e. surrogate substrates) concept | [43] |
| FFN54/246 | SLC6A4, SLC18 | 'False' fluorescent substrates for serotonin and VMAT transporters. Potent inhibition by imipramine and citalopram | [44] |
| FFN270 | SLC6A4, SLC18 | Another example of a fluorescent false neurotransmitter | [119] |
| Fluorescein | SLCO1B1/3B1 | Effective substrate; analysis of inhibitors | [58] [59] |
|  | OAT6 (SLC22A20) |  | [60] |
|  | SLC16A1, SLC16A4 |  | [57] |
|  | Many OATPs (SLCO family) expressed in insect cells |  | [56] |
| Lucifer yellow | Sodium-dependent anion transporters | Inhibited by probenecid | [120; 121] |
| Rhodamine 123 | OCT1/OCT2 (SLC22A1/2) | Potent substrate | [122] |
| Stilbazolium dyes | Norepinephrine transporter (SLC6) | Dyes related to ASP<sup>+</sup> | [123] |
| Zombie Violet, Live/Dead Green, Cascade Blue, Alexa Fluor 405 | OATP (SLCO) 1B1/1B3 and 2B1 | All shown to be direct substrates, and uptake inhibited by known transporter inhibitors | [124] |

Fluorescein is similar in t-SNE space (Fig 4A) to a variety of drugs. This similarity is not at all related to the class of drug, however, as close ones include balsalazide (an anti-inflammatory used in inflammatory bowel disease [61]), bentiromide (a peptide used for assessing pancreatic function [62]), butenafine (a topical antifungal [63]), sertindole (an atypical antipsychotic), and tolvaptan (used in autosomal dominant polycystic kidney disease [64]). Similar remarks may be made of dapoxyl (Fig 4B). Note, of course, that the t-SNE plots are based on property descriptors, while the Tanimoto distances are based on a particular form of molecular fingerprint, so *a priori* we do not necessarily expect the closest molecules to be the same in the two cases. In addition, we note that molecules with different scaffolds may be quite similar; in the cheminformatics literature this is known as ‘scaffold hopping’ (e.g. [65–70]).

For a drug, we picked nitisinone, a drug active against hereditary tyrosinaemia type I [71] and alkaptonuria [72; 73], as it is surrounded in t-SNE space (Fig 4C) by several tricyclic fluorophores, that do indeed share similar structures (Fig 4C).

Bickerton and colleagues [74] introduced the concept of the quantitative estimate of drug-likeness (QED) (but see [75]), and it is of interest to see how ‘drug-like’ our four classes of molecule are by their criteria. Fig 5A shows the distribution of QED drug-likenesses for marketed drugs, for Recon2 metabolites, for our selected fluorophores, and for a sample of 2000 molecules from UNPD. Our fluorophores are noticeably more similar to drugs than are endogenous metabolites, and roughly as similar to drugs as are natural products (Fig 5A).

**Figure 5.**
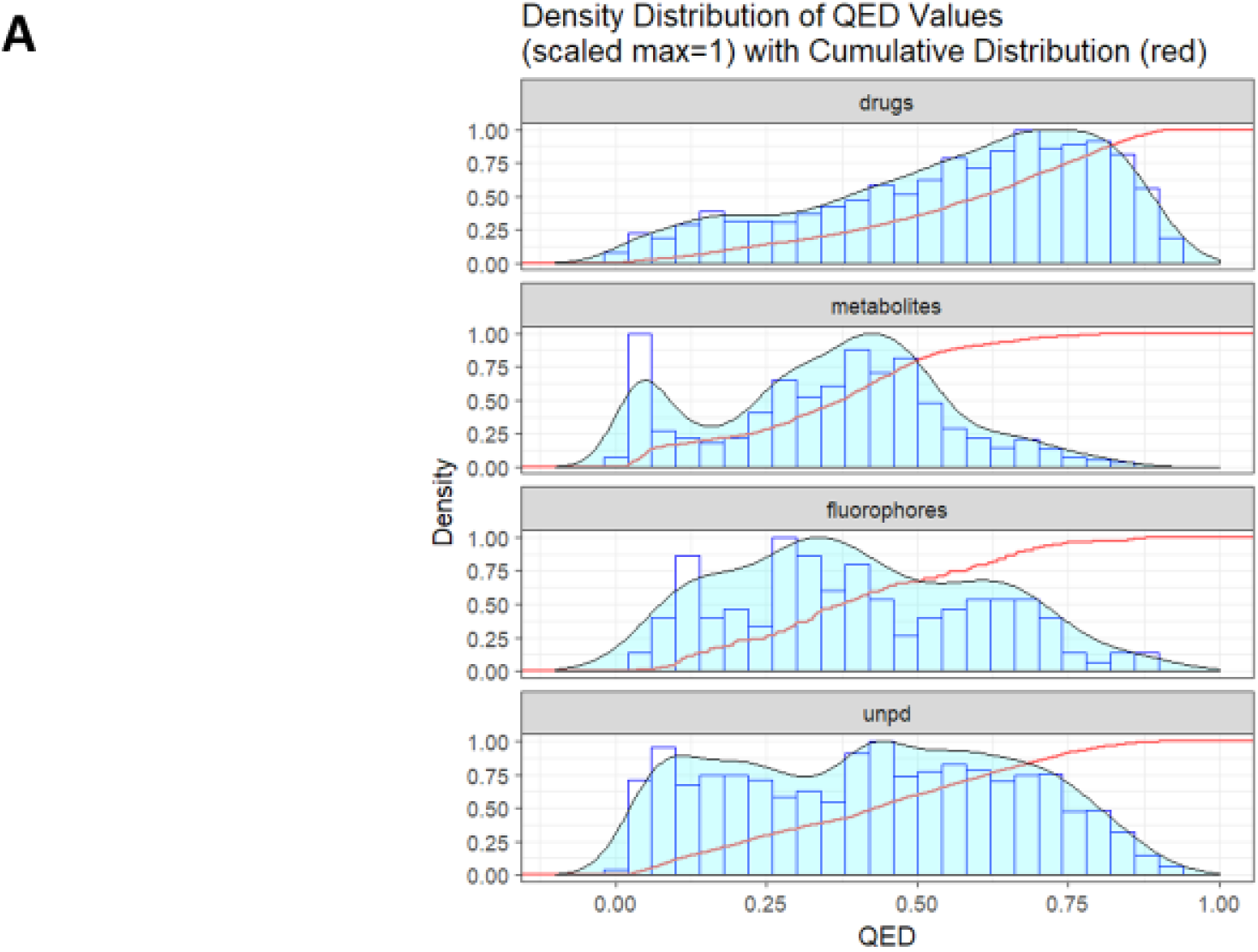
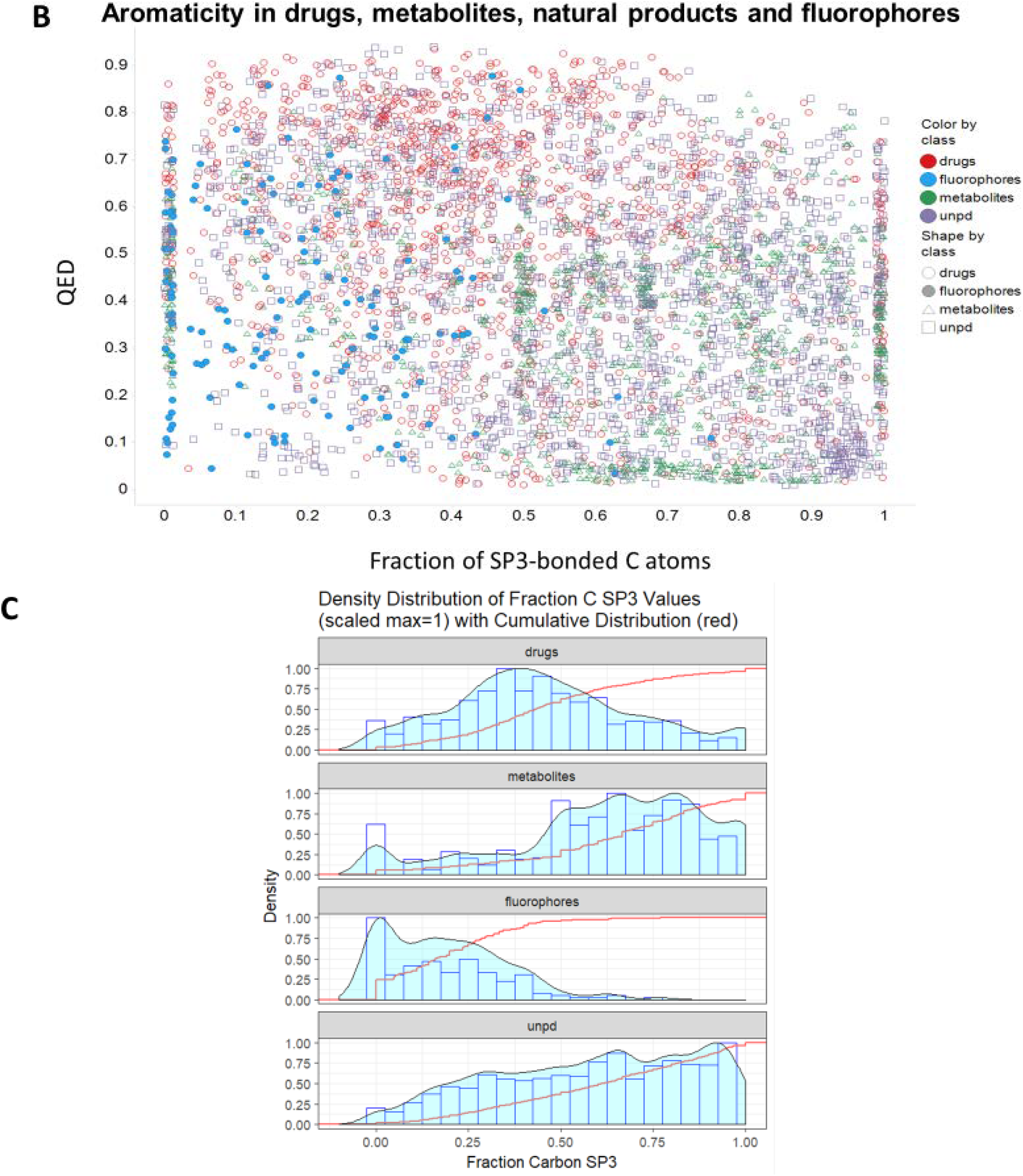
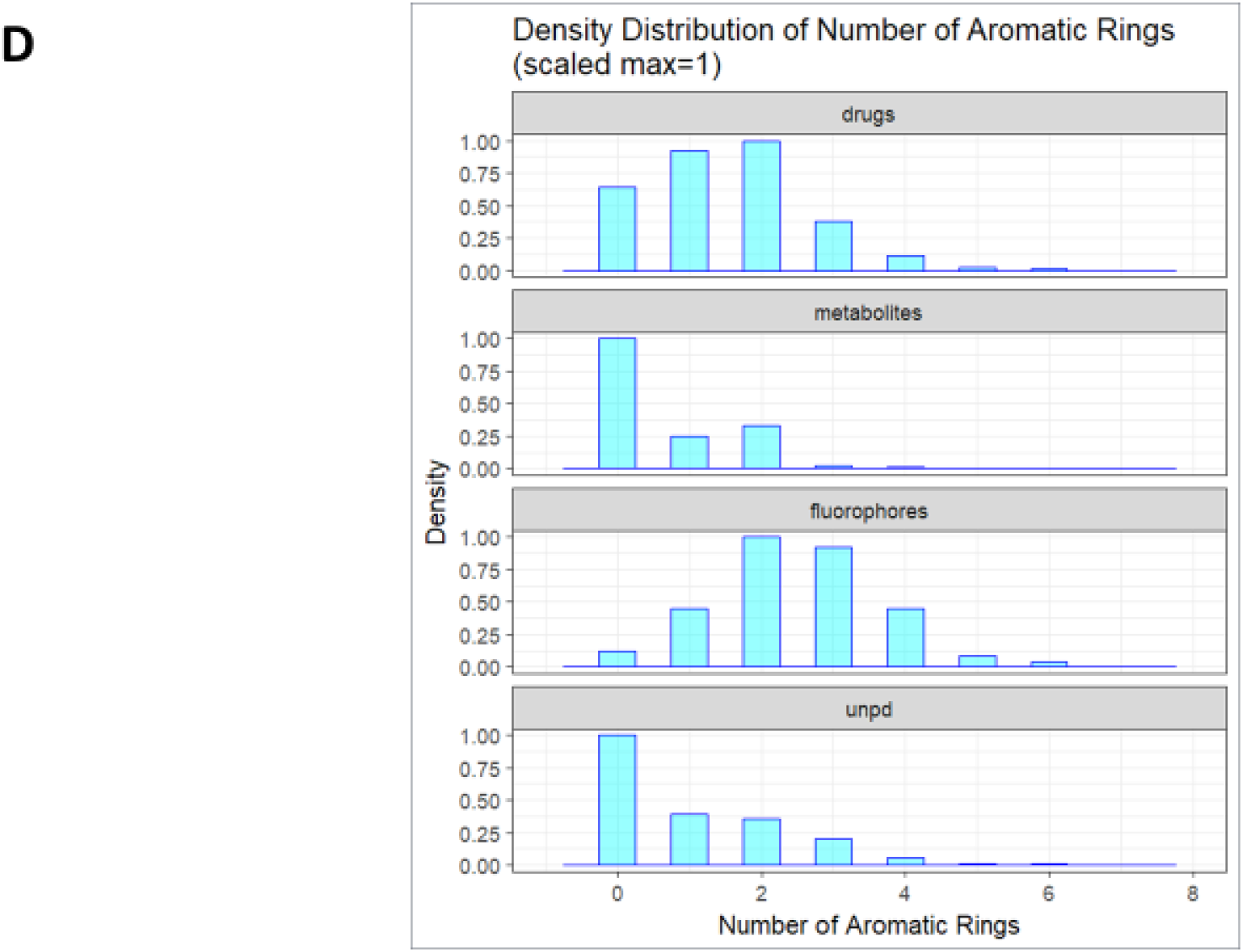
Distribution of quantitative estimate of drug-likeness (QED) values in different classes of molecule. A. Cumulative distributions for the four classes. B. Relationship between QED and aromaticity for the four classes as encoded by the fraction of C atoms exhibiting sp^3^ bonding. QED values were calculated using the RDKit Python code as described in Methods and plotted in (A) using ggplot2 and in (B) using Spotfire. C. Density distribution of fraction of C atoms with sp^3^ bonding. D. Histogram of distributions of numbers of aromatic rings in the four given classes.

Given that essentially all drugs are similar to at least one natural product [42], this is entirely consistent with our thesis that most fluorophores do look rather like one or more marketed drugs. One aspect in which (a) drugs and fluorophores differ noticeably from (b) metabolites and natural products is the extent to which they exhibit aromaticity, here encoded (Fig 5B, on the abscissa) via the fraction of carbon atoms showing sp^3^ hybridisation (i.e. non-aromatic). This is shown as a distribution in Fig 5C. There is clearly a significant tendency for drugs to include (planar) aromatic rings, and although this is changing somewhat [76–80] there are strong thermodynamic reasons as to why this should be so (see Discussion). The modal number of aromatic rings for both drugs and fluorophores is two, significantly greater than that (zero) for metabolites and for natural products (Fig 5D). One reason for fluorophores to exhibit aromaticity is simple, as reasonable visible-wavelength fluorescence in organic molecules relies strongly on conjugation (e.g. [81]), to which aromatic rings can contribute strongly. This argument alone probably accounts in large measure for the drug-likeness of fluorophores.

Finally, a very recent, principled, and effective clustering method [82; 83], representing the state of the art, is that based on the Uniform Manifold Approximation and Projection (UMAP) algorithm. In a similar vein, and based on the same descriptors as used in the t-SNE plots, we show the clustering of our four classes of molecule in UMAP space, where most clusters containing drugs also contain fluorophores. Despite being based on property descriptors, the UMAP algorithm is clearly very effective at clustering molecules into structurally related classes.

## DISCUSSION

The basis of the main idea presented here is that the structures of common fluorophores are sufficiently similar to those of many drugs as to provide suitable surrogates for assessing their uptake via solute carriers of the SLC (and indeed their efflux via ABC) families. While the latter transporters are well known to be rather promiscuous, and to transport a variety of fluorophores [34; 36; 84-86], considerably less attention has been paid to the former. Of course some marketed pharmaceutical drugs that are transported into cells are in fact naturally fluorescent, including molecules such as anthracyclines [87–89], mepacrine (atebrin, quinacrine) [90], obatoclax [91; 92], tetracycline derivatives [88; 93] and topotecan [94], The same is true of certain vitamins such as riboflavin [95; 96] (that necessarily have transporters, as cells cannot synthesise them), as well as certain bioactive natural products (e.g. [97–99]). As an illustration, and as a complement to our detailed gene knockout studies [17], Table 1 gives an indication of dyes whose interaction with specific transporters has been demonstrated directly. In some cases, their surrogacy as a substrate for a transporter with a known non-fluorescent substrate is clear, and as mentioned in the introduction they are sometimes referred to as ‘false fluorescent substrates’. Overall, while not intended to be remotely exhaustive, this Table does serve to indicate the potentially widespread activity of transporters as mediators of fluorophore uptake, and indeed a number of such transporters are known to be rather promiscuous.

Structural similarity (or the assessment of properties based simply on analyzing structures) is an elusive concept (e.g. [125]), but as judged by a standard encoding (RDKit patterned) there is considerable similarity in structure between almost all of our chosen fluorophores and at least one drug, whether this is judged by their descriptor- or fingerprint-based properties (Figs 1-3), by observation (Figs 4, 6), or (Fig 5) via the QED [74] measure.

**Fig 6.**
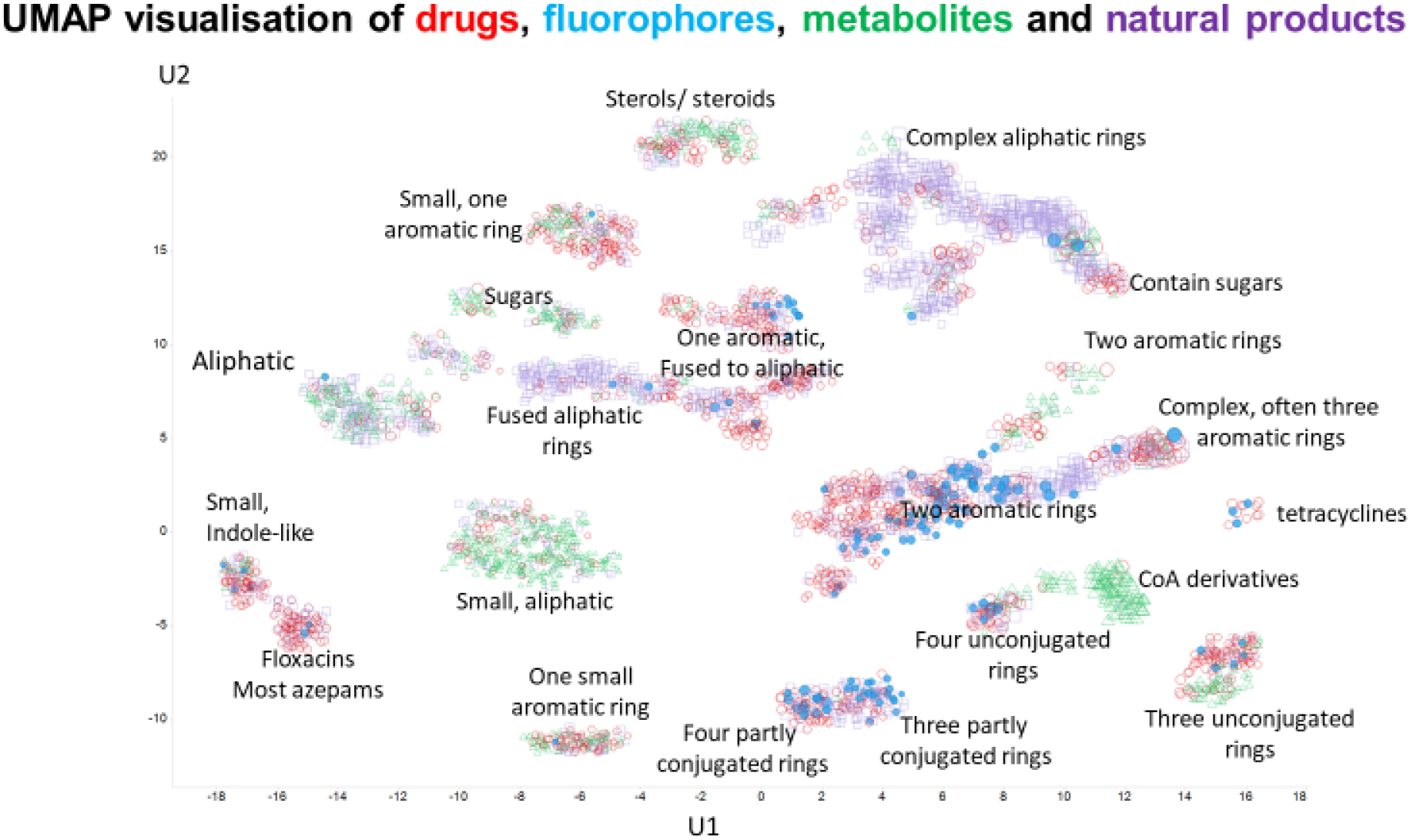
UMAP projection into two dimensions of the four classes of molecules, annotated by the type of molecular structure in the various clusters.

Although there is a move to phenotypic screening [126–129], many drugs were developed on the basis of their ability to bind potently *in vitro* to a target of interest. If the unbound molecule is conformationally very flexible, and the bound version is not, binding necessarily involves a significant loss of entropy. Potent binding (involving a significant loss in free energy) of such a molecule would thus require a very large enthalpic term. Consequently, it is much easier to find potent binders if the binding can involve flat (which implies aromatic), conformationally inflexible planar structures. Such reasoning presumably reflects the observation (Fig 5B) that drugs tend to have a low sp^3^ character, typically with a number of aromatic rings. Conjugated aromatic rings are also a major (physical and electronic) structure that allow fluorescence from organic molecules [130–133], with greater π-bond conjugation moving both absorbance and fluorescence toward the red end of the spectrum. Overall, these two separate roles for aromatic residues, in low entropy of binding and in electronic structure, provide a plausible explanation for much of the drug-likeness of common fluorophores.

While this study used a comparatively small set of fluorophores, increasing their number can only increase the likelihood of finding a drug (or natural product) to which they are seen to be similar. This said, this set of molecules provides an excellent starting point for the development of competitive high-throughput assays of drug transporter activity.

## CONCLUSIONS

An analysis of some 150 fluorophores in common usage in biological research has shown that a very great many of them bear significant structural similarities to marketed drugs (and to natural products). This similarity holds true whether the analysis is done using structures encoded as fingerprints or via physico-chemical descriptors, by visual inspection, or via the quantitative estimate of drug likeness measure. For any given drug there is thus likely to be a fluorophore or set of fluorophores that is best suited to competing with it for uptake, and thus for determining by fluorimetric methods the QSAR for the relevant transporters. This should provide the means for rapid and convenient competitive and trans-stimulation assays for screening the ability of drugs to enter cells via SLCs.

## ABBREVIATIONS

PCA: Principal Components Analysis
QED: Quantitative Estimate of Drug-likeness
QSAR: Quantittaive Structure-Activity Relationship
SLC: solute carrier
TS: Tanimoto similarity
UNPD: Universal Natural Products Database

## SUPPLEMENTARY MATERIALS

The following supplementary materials are available online in the file fluorophoresSI.xIsx. Table S1: The list of all the molecules and properties used in the present analysis.

## DATA AVAILABILITY

- The dataset generated from (or analyzed in) the study can be found in the Supplementary Excel sheet entitled fluorophoresSI.xlsx.

## AUTHOR CONTRIBUTIONS

DBK developed the idea of determining fluorophore-drug similarity. SO’H wrote and ran the majority of the workflows and created ~two-thirds of the visualisations. Both authors contributed to the analysis of the data and to the writing of the paper.

## How to cite this article

O’Hagan S, Kell DB. Structural similarities between some common fluorophores used in biology and marketed drugs, endogenous metabolites, and natural products. Pharm Front. 2019, submitted for review.

## REFERENCES

[1] Chalfie, M. & Kain, S. (1998). Green Fluorescent Protein: properties, applications, and protocols. Wiley-Liss, New York.

[2] Hemmilä, I. A. (1991). Applications of fluorescence in immunoassays. Wiley, New York.

[3] Waggoner, A. S. (1990). Fluorescent Probes for Cytometry. In Flow Cytometry and Sorting (2nd Edition) (ed. M. R. Melamed, T. Lindmo and M. L. Mendelsohn), pp. 209–225. Wiley-Liss Inc., New York.

[4] Kyrychenko, A. (2015). Using fluorescence for studies of biological membranes: a review. Methods Appl Fluoresc 3, 042003.

[5] Lagorio, M. G., Cordon, G. B. & Iriel, A. (2015). Reviewing the relevance of fluorescence in biological systems. Photochem Photobiol Sci 14, 1538–59.

[6] Specht, E. A., Braselmann, E. & Palmer, A. E. (2017). A Critical and Comparative Review of Fluorescent Tools for Live-Cell Imaging. Annu Rev Physiol 79, 93–117.

[7] Sahl, S. J., Hell, S. W. & Jakobs, S. (2017). Fluorescence nanoscopy in cell biology. Nat Rev Mol Cell Biol 18, 685–701.

[8] Mavrakis, M., Pourquie, O. & Lecuit, T. (2010). Lighting up developmental mechanisms: how fluorescence imaging heralded a new era. Development 137, 373–87.

[9] Johnson, I. (1998). Fluorescent probes for living cells. Histochem J 30, 123–40.

[10] Kolanowski, J. L., Liu, F. & New, E. J. (2018). Fluorescent probes for the simultaneous detection of multiple analytes in biology. Chem Soc Rev 47, 195–208.

[11] Zhang, J., Campbell, R. E., Ting, A. Y. & Tsien, R. Y. (2002). Creating new fluorescent probes for cell biology. Nat Rev Mol Cell Biol 3, 906–18.

[12] Jiang, X., Wang, L., Carroll, S. L., Chen, J., Wang, M. C. & Wang, J. (2018). Challenges and Opportunities for Small-Molecule Fluorescent Probes in Redox Biology Applications. Antioxid Redox Signal 29, 518–540.

[13] Winterbourn, C. C. (2014). The challenges of using fluorescent probes to detect and quantify specific reactive oxygen species in living cells. Biochim Biophys Acta 1840, 730–8.

[14] Davey, H. M. & Kell, D. B. (1996). Flow cytometry and cell sorting of heterogeneous microbial populations: the importance of single-cell analysis. Microbiol. Rev. 60, 641–696.

[15] Shapiro, H. M. (2003). Practical Flow Cytometry, 4th edition, 3rd edition. John Wiley, New York.

[16] Lavis, L. D. & Raines, R. T. (2008). Bright ideas for chemical biology. ACS Chem Biol 3, 142–55.

[17] Jindal, S., Yang, L., Day, P. J. & Kell, D. B. (2019). Involvement of multiple influx and efflux transporters in the accumulation of cationic fluorescent dyes by *Escherichia coli*. BMC Microbiol 19, 195; also bioRxiv 603688v1.

[18] Kell, D. B. (2019). Control of metabolite efflux in microbial cell factories: current advances and future prospects. In Fermentation microbiology and biotechnology, 4th Ed (ed. E. M. T. El-Mansi, J. Nielsen, D. Mousdale, T. Allman and R. Carlson), pp. 117–138. CRC Press, Boca Raton.

[19] Szakács, G., Váradi, A., Özvegy-Laczka, C. & Sarkadi, B. (2008). The role of ABC transporters in drug absorption, distribution, metabolism, excretion and toxicity (ADME-Tox). Drug Discov Today 13, 379–93.

[20] Schinkel, A. H. & Jonker, J. W. (2012). Mammalian drug efflux transporters of the ATP binding cassette (ABC) family: an overview. Adv Drug Deliv Rev 64, 138–153.

[21] Tar ling, E. J., de Aguiar Vallim, T. Q. & Edwards, P. A. (2013). Role of ABC transporters in lipid transport and human disease. Trends Endocrinol Metab 24, 342–350.

[22] Andreoletti, P., Raas, Q., Gondcaille, C., Cherkaoui-Malki, M., Trompier, D. & Savary, S. (2017). Predictive Structure and Topology of Peroxisomal ATP-Binding Cassette (ABC) Transporters. Int J Mol Sci 18.

[23] Sharom, F. J. (2008). ABC multidrug transporters: structure, function and role in chemoresistance. Pharmacogenomics 9, 105–27.

[24] Pahnke, J., Fröhlich, C., Paarmann, K., Krohn, M., Bogdanovic, N., Årsland, D. & Winblad, B. (2014). Cerebral ABC transporter-common mechanisms may modulate neurodegenerative diseases and depression in elderly subjects. Arch Med Res 45, 738–43.

[25] Abuznait, A. H. & Kaddoumi, A. (2012). Role of ABC transporters in the pathogenesis of Alzheimer’s disease. ACS Chem Neurosci 3, 820–31.

[26] Lam, F. C., Liu, R., Lu, P., Shapiro, A. B., Renoir, J. M., Sharom, F. J. & Reiner, P. B. (2001). beta-Amyloid efflux mediated by p-glycoprotein. J Neurochem 76, 1121–8.

[27] Molnar, J., Ocsovszki, I. & Pusztai, R. (2018). Amyloid-beta Interactions with ABC Transporters and Resistance Modifiers. Anticancer Res 38, 3407–3410.

[28] Wijnholds, J., Evers, R., van Leusden, M. R., Mol, C. A. A. M., Zaman, G. J. R., Mayer, U., Beijnen, J. H., van der Valk, M., Krimpenfort, P. & Borst, P. (1997). Increased sensitivity to anticancer drugs and decreased inflammatory response in mice lacking the multidrug resistance-associated protein. Nat Med 3, 1275–9.

[29] Prates, R. A., Kato, I. T., Ribeiro, M. S., Tegos, G. P. & Hamblin, M. R. (2011). Influence of multidrug efflux systems on methylene blue-mediated photodynamic inactivation of Candida albicans. J Antimicrob Chemother 66, 1525–32.

[30] Forster, S., Thumser, A. E., Hood, S. R. & Plant, N. (2012). Characterization of rhodamine-123 as a tracer dye for use in *in vitro drug* transport assays. PLoS One 7, e33253.

[31] Ivnitski-Steele, I., Holmes, A. R., Lamping, E., Monk, B. C., Cannon, R. D. & Sklar, L. A. (2009). Identification of Nile red as a fluorescent substrate of the *Candida albicans* ATP-binding cassette transporters Cdr1p and Cdr2p and the major facilitator superfamily transporter Mdr1p. Anal Biochem 394, 87–91.

[32] Strouse, J. J., Ivnitski-Steele, I., Waller, A., Young, S. M., Perez, D., Evangelisti, A. M., Ursu, O., Bologa, C. G., Carter, M. B., Salas, V. M., Tegos, G., Larson, R. S., Oprea, T. I., Edwards, B. S. & Sklar, L. A. (2013). Fluorescent substrates for flow cytometric evaluation of efflux inhibition in ABCB1, ABCC1, and ABCG2 transporters. Anal Biochem 437, 77–87.

[33] Kell, D. B. (2018). Control of metabolite efflux in microbial cell factories: current advances and future prospects. OSF preprints, xg9jh.

[34] Szabó, E., Türk, D., Telbisz, Á., Kucsma, N., Horváth, T., Szakács, G., Homolya, L., Sarkadi, B. & Várady, G. (2018). A new fluorescent dye accumulation assay for parallel measurements of the ABCG2, ABCB1 and ABCC1 multidrug transporter functions. PLoS One 13, e0190629.

[35] Gökirmak, T., Shipp, L. E., Campanale, J. P., Nicklisch, S. C. T. & Hamdoun, A. (2014). Transport in Technicolor: Mapping ATP-Binding Cassette Transporters in Sea Urchin Embryos. Mol Reprod Dev 81, 778–793.

[36] Fardel, O., Le Vee, M., Jouan, E., Denizot, C. & Parmentier, Y. (2015). Nature and uses of fluorescent dyes for drug transporter studies. Expert Opin Drug Metab Toxicol 11, 1233–1251.

[37] O’Hagan, S., Swainston, N., Handl, J. & Kell, D. B. (2015). A ‘rule of 0.5’ for the metabolite-likeness of approved pharmaceutical drugs. Metabolomics 11, 323–339.

[38] O’Hagan, S. & Kell, D. B. (2015). Understanding the foundations of the structural similarities between marketed drugs and endogenous human metabolites. Front Pharmacol 6, 105.

[39] O’Hagan, S. & Kell, D. B. (2016). MetMaxStruct: a Tversky-similarity-based strategy for analysing the (sub)structural similarities of drugs and endogenous metabolites. Front Pharmacol 7, 266.

[40] O’Hagan, S. & Kell, D. B. (2017). Analysis of drug-endogenous human metabolite similarities in terms of their maximum common substructures. J Cheminform 9, 18.

[41] Gründemann, D., Harlfinger, S., Golz, S., Geerts, A., Lazar, A., Berkels, R., Jung, N., Rubbert, A. & Schömig, E. (2005). Discovery of the ergothioneine transporter. Proc Natl Acad Sci 102, 5256–61.

[42] O’Hagan, S. & Kell, D. B. (2017). Consensus rank orderings of molecular fingerprints illustrate the ‘most genuine’ similarities between marketed drugs and small endogenous human metabolites, but highlight exogenous natural products as the most important ‘natural’ drug transporter substrates. ADMET & DMPK 5, 85–125.

[43] Gubernator, N. G., Zhang, H., Staal, R. G., Mosharov, E. V., Pereira, D. B., Yue, M., Balsanek, V., Vadola, P. A., Mukherjee, B., Edwards, R. H., Sulzer, D. & Sames, D. (2009). Fluorescent false neurotransmitters visualize dopamine release from individual presynaptic terminals. Science 324, 1441–4.

[44] Henke, A., Kovalyova, Y., Dunn, M., Dreier, D., Gubernator, N. G., Dincheva, I., Hwu, C., Sebej, P., Ansorge, M. S., Sulzer, D. & Sames, D. (2018). Toward Serotonin Fluorescent False Neurotransmitters: Development of Fluorescent Dual Serotonin and Vesicular Monoamine Transporter Substrates for Visualizing Serotonin Neurons. ACS Chem Neurosci 9, 925–934.

[45] Thiele, I., Swainston, N., Fleming, R. M. T., Hoppe, A., Sahoo, S., Aurich, M. K., Haraldsdottír, H., Mo, M. L., Rolfsson, O., Stobbe, M. D., Thorleifsson, S. G., Agren, R., Bölling, C., Bordel, S., Chavali, A. K., Dobson, P., Dunn, W. B., Endler, L., Goryanin, I., Hala, D., Hucka, M., Hull, D., Jameson, D., Jamshidi, N., Jones, J., Jonsson, J. J., Juty, N., Keating, S., Nookaew, I., Le Novère, N., Malys, N., Mazein, A., Papin, J. A., Patel, Y., Price, N. D., Selkov Sr., E., Sigurdsson, M. I., Simeonidis, E., Sonnenschein, N., Smallbone, K., Sorokin, A., Beek, H. V., Weichart, D., Nielsen, J. B., Westerhoff, H. V., Kell, D. B., Mendes, P. & Palsson, B. Ø. (2013). A community-driven global reconstruction of human metabolism. Nat Biotechnol. 31, 419–425.

[46] Gu, J. Y., Gui, Y. S., Chen, L. R., Yuan, G., Lu, H. Z. & Xu, X. J. (2013). Use of Natural Products as Chemical Library for Drug Discovery and Network Pharmacology. PloS one 8, e62839.

[47] O’Hagan, S. & Kell, D. B. (2015). The KNIME workflow environment and its applications in Genetic Programming and machine learning. Genetic Progr Evol Mach 16, 387–391.

[48] O’Hagan, S. & Kell, D. B. (2015). The apparent permeabilities of Caco-2 cells to marketed drugs: magnitude, and independence from both biophysical properties and endogenite similarities Peer J 3, e1405.

[49] O’Hagan, S. & Kell, D. B. (2018). Analysing and navigating natural products space for generating small, diverse, but representative chemical libraries. Biotechnol J 13, 1700503.

[50] O’Hagan, S. & Kell, D. B. (2019). Generation of a small library of natural products designed to cover chemical space inexpensively. Pharm Front 1, e190005.

[51] van der Maaten, L. & Hinton, G. (2008). Visualizing Data using t-SNE. J Machine Learning Res 9, 2579–2605.

[52] Thiele, I., Heinken, A. & Fleming, R. M. (2013). A systems biology approach to studying the role of microbes in human health. Curr Opin Biotechnol 24, 4–12.

[53] Eisen, M. B., Spellman, P. T., Brown, P. O. & Botstein, D. (1998). Cluster analysis and display of genome-wide expression patterns. Proc. Natl. Acad. Sci. 95, 14863–14868.

[54] Bakkar, M. M., Hardaker, L., March, P., Morgan, P. B., Maldonado-Codina, C. & Dobson, C. B. (2014). The cellular basis for biocide-induced fluorescein hyperfluorescence in mammalian cell culture. PLoS One 9, e84427.

[55] Khan, T. F., Price, B. L., Morgan, P. B., Maldonado-Codina, C. & Dobson, C. B. (2018). Cellular fluorescein hyperfluorescence is dynamin-dependent and increased by Tetronic 1107 treatment. Int J Biochem Cell Biol 101, 54–63.

[56] Patik, I., Kovacsics, D., Német, O., Gera, M., Várady, G., Stieger, B., Hagenbuch, B., Szakács, G. & Özvegy-Laczka, C. (2015). Functional expression of the 11 human Organic Anion Transporting Polypeptides in insect cells reveals that sodium fluorescein is a general OATP substrate. Biochem Pharmacol 98, 649–58.

[57] Sun, Y. C., Liou, H. M., Yeh, P. T., Chen, W. L. & Hu, F. R. (2017). Monocarboxylate Transporters Mediate Fluorescein Uptake in Corneal Epithelial Cells. Invest Ophthalmol Vis Sci 58, 3716–3722.

[58] De Bruyn, T., Fattah, S., Stieger, B., Augustijns, P. & Annaert, P. (2011). Sodium fluorescein is a probe substrate for hepatic drug transport mediated by OATP1B1 and OATP1B3. J Pharm Sci 100, 5018–30.

[59] De Bruyn, T., van Westen, G. J., Ijzerman, A. P., Stieger, B., de Witte, P., Augustijns, P. F. & Annaert, P. P. (2013). Structure-based identification of OATP1B1/3 inhibitors. Mol Pharmacol 83, 1257–67.

[60] Truong, D. M., Kaler, G., Khandelwal, A., Swaan, P. W. & Nigam, S. K. (2008). Multi-level analysis of organic anion transporters 1, 3, and 6 reveals major differences in structural determinants of antiviral discrimination. J Biol Chem 283, 8654–63.

[61] Patil, S. A. & Moss, A. C. (2008). Balsalazide disodium for the treatment of ulcerative colitis. Expert Rev Gastroenterol Hepatol 2, 177–84.

[62] Egesel, T., Ünsal, I., Dikmen, G. & Bayraktar, Y. (2002). The assessment of pancreatic exocrine function by bentiromide test in patients with chronic portal vein thrombosis. Pancreas 25, 355–9.

[63] Singal, A. (2008). Butenafine and superficial mycoses: current status. Expert Opin Drug Metab Toxicol 4, 999–1005.

[64] Blair, H. A. (2019). Tolvaptan: A Review in Autosomal Dominant Polycystic Kidney Disease. Drugs 79, 303–313.

[65] Brown, N. & Jacoby, E. (2006). On scaffolds and hopping in medicinal chemistry. Mini Rev Med Chem 6, 1217–29.

[66] Geppert, H. & Bajorath, J. (2010). Advances in 2D fingerprint similarity searching. Expert Opin Drug Discov 5, 529–542.

[67] Lamberth, C. (2018). Agrochemical lead optimization by scaffold hopping. Pest Manag Sci 74, 282–292.

[68] Mauser, H. & Guba, W. (2008). Recent developments in *de novo* design and scaffold hopping. Curr Opin Drug Discov Devel 11, 365–374.

[69] Sun, H., Tawa, G. & Wallqvist, A. (2012). Classification of scaffold-hopping approaches. Drug Discov Today 17, 310–24.

[70] Zhao, H. (2007). Scaffold selection and scaffold hopping in lead generation: a medicinal chemistry perspective. Drug Disc Today 12, 149–55.

[71] Das, A. M. (2017). Clinical utility of nitisinone for the treatment of hereditary tyrosinemia type-1 (HT-1). Appl Clin Genet 10, 43–48.

[72] Lock, E., Ranganath, L. R. & Timmis, O. (2014). The role of nitisinone in tyrosine pathway disorders. Curr Rheumatol Rep 16, 457.

[73] Ranganath, L. R., Khedr, M., Milan, A. M., Davison, A. S., Hughes, A. T., Usher, J. L., Taylor, S., Loftus, N., Daroszewska, A., West, E., Jones, A., Briggs, M., Fisher, M., McCormick, M., Judd, S., Vinjamuri, S., Griffin, R., Psarelli, E. E., Cox, T. F., Sireau, N., Dillon, J. P., Devine, J. M., Hughes, G., Harrold, J., Barton, G. J., Jarvis, J. C. & Gallagher, J. A. (2018). Nitisinone arrests ochronosis and decreases rate of progression of Alkaptonuria: Evaluation of the effect of nitisinone in the United Kingdom National Alkaptonuria Centre. Mol Genet Metab 125, 127–134.

[74] Bickerton, G. R., Paolini, G. V., Besnard, J., Muresan, S. & Hopkins, A. L. (2012). Quantifying the chemical beauty of drugs. Nat Chem 4, 90–8.

[75] Shultz, M. D. (2019). Two Decades under the Influence of the Rule of Five and the Changing Properties of Approved Oral Drugs. J Med Chem 62, 1701–1714.

[76] Lovering, F., Bikker, J. & Humblet, C. (2009). Escape from flatland: increasing saturation as an approach to improving clinical success. J Med Chem 52, 6752–6.

[77] Meyers, J., Carter, M., Mok, N. Y. & Brown, N. (2016). On the origins of three-dimensionality in drug-like molecules. Future Med Chem 8, 1753–67.

[78] Campbell, P. S., Jamieson, C., Simpson, I. & Watson, A. J. B. (2017). Practical synthesis of pharmaceutically relevant molecules enriched in sp(3) character. Chem Commun (Camb) 54, 46–49.

[79] Blakemore, D. C., Castro, L., Churcher, I., Rees, D. C., Thomas, A. W., Wilson, D. M. & Wood, A. (2018). Organic synthesis provides opportunities to transform drug discovery. Nat Chem 10, 383–394.

[80] Boström, J., Brown, D. G., Young, R. J. & Keserü, G. M. (2018). Expanding the medicinal chemistry synthetic toolbox. Nat Rev Drug Discov 17, 709–727.

[81] Yuan, L., Lin, W., Zheng, K., He, L. & Huang, W. (2013). Far-red to near infrared analyte-responsive fluorescent probes based on organic fluorophore platforms for fluorescence imaging. Chem Soc Rev 42, 622–61.

[82] McInnes, L., Healy, J. & Melville, J. (2018). UMAP: Uniform Manifold Approximation and Projection for Dimension Reduction. arXiv, 1802.03426v2.

[83] McInnes, L., Healy, J., Saul, N. & Großberger, L. (2018). UMAP: Uniform Manifold Approximation and Projection. J Open Source Software DOI 10.21105/joss.00861.

[84] Ivnitski-Steele, I., Larson, R. S., Lovato, D. M., Khawaja, H. M., Winter, S. S., Oprea, T. I., Sklar, L. A. & Edwards, B. S. (2008). High-throughput flow cytometry to detect selective inhibitors of ABCB1, ABCC1, and ABCG2 transporters. Assay Drug Dev Technol 6, 263–76.

[85] Tegos, G. P., Evangelisti, A. M., Strouse, J. J., Ursu, O., Bologa, C. & Sklar, L. A. (2014). A high throughput flow cytometric assay platform targeting transporter inhibition. Drug Disc Today Technol 12, e95–e103.

[86] Windt, T., Tóth, S., Patik, I., Sessler, J., Kucsma, N., Szepesi, A., Zdrazil, B., Özvegy-Laczka, C. & Szakács, G. (2019). Identification of anticancer OATP2B1 substrates by an in vitro triple-fluorescence-based cytotoxicity screen. Arch Toxicol 93, 953–964.

[87] Gautier, J., Munnier, E., Souce, M., Chourpa, I. & Douziech Eyrolles, L. (2015). Analysis of doxorubicin distribution in MCF-7 cells treated with drug-loaded nanoparticles by combination of two fluorescence-based techniques, confocal spectral imaging and capillary electrophoresis. Anal Bioanal Chem.

[88] Khader, H., Solodushko, V., Al-Mehdi, A. B., Audia, J. & Fouty, B. (2014). Overlap of Doxycycline Fluorescence with that of the Redox-Sensitive Intracellular Reporter roGFP. Journal of Fluorescence 24, 305–311.

[89] Motlagh, N. S. H., Parvin, P., Ghasemi, F. & Atyabi, F. (2016). Fluorescence properties of several chemotherapy drugs: doxorubicin, paclitaxel and bleomycin. Biomed Opt Express 7, 2400–6.

[90] Baldini, G., Doglia, S., Dolci, S. & Sassi, G. (1981). Fluorescence-determined preferential binding of quinacrine to DNA. Biophys J 36, 465–77.

[91] Nguyen, M., Marcellus, R. C., Roulston, A., Watson, M., Serfass, L., Murthy Madiraju, S. R., Goulet, D., Viallet, J., Belec, L., Billot, X., Acoca, S., Purisima, E., Wiegmans, A., Cluse, L., Johnstone, R. W., Beauparlant, P. & Shore, G. C. (2007). Small molecule obatoclax (GX15-070) antagonizes MCL-1 and overcomes MCL-1-mediated resistance to apoptosis. Proc Natl Acad Sci U S A 104, 19512–7.

[92] Stamelos, V. A., Fisher, N., Bamrah, H., Voisey, C., Price, J. C., Farrell, W. E., Redman, C. W. & Richardson, A. (2016). The BH3 Mimetic Obatoclax Accumulates in Lysosomes and Causes Their Alkalinization. PLoS One 11, e0150696.

[93] Pautke, C., Vogt, S., Kreutzer, K., Haczek, C., Wexel, G., Kolk, A., Imhoff, A. B., Zitzelsberger, H., Milz, S. & Tischer, T. (2010). Characterization of eight different tetracyclines: advances in fluorescence bone labeling. J Anat 217, 76–82.

[94] Burke, T. G., Malak, H., Gryczynski, I., Mi, Z. & Lakowicz, J. R. (1996). Fluorescence detection of the anticancer drug topotecan in plasma and whole blood by two-photon excitation. Anal Biochem 242, 266–70.

[95] Yonezawa, A. & Inui, K. (2013). Novel riboflavin transporter family RFVT/SLC52: identification, nomenclature, functional characterization and genetic diseases of RFVT/SLC52. Mol Aspects Med 34, 693–701.

[96] Zhang, S., Sakuma, M., Deora, G. S., Levy, C. W., Klausing, A., Breda, C., Read, K. D., Edlin, C. D., Ross, B. P., Wright Muelas, M., Day, P. J., O’Hagan, S., Kell, D. B., Schwarcz, R., Leys, D., Heyes, D. J., Giorgini, F. & Scrutton, N. S. (2019). A brain-permeable inhibitor of the neurodegenerative disease target kynurenine 3-monooxygenase prevents accumulation of neurotoxic metabolites. Commun Biol 2, 271.

[97] Liu, Q., Liu, Y., Guo, M., Luo, X. & Yao, S. (2006). A simple and sensitive method of nonaqueous capillary electrophoresis with laser-induced native fluorescence detection for the analysis of chelerythrine and sanguinarine in Chinese herbal medicines. Talanta 70, 202–7.

[98] Duval, R. & Duplais, C. (2017). Fluorescent natural products as probes and tracers in biology. Nat Prod Rep 34, 161–193.

[99] Taniguchi, M. & Lindsey, J. S. (2018). Database of Absorption and Fluorescence Spectra of >300 Common Compounds for use in PhotochemCAD. Photochem Photobiol 94, 290–327.

[100] Bednarczyk, D., Mash, E. A., Aavula, B. R. & Wright, S. H. (2000). NBD-TMA: a novel fluorescent substrate of the peritubular organic cation transporter of renal proximal tubules. Pflugers Arch 440, 184–92.

[101] Stone, M. R. L., Butler, M. S., Phetsang, W., Cooper, M. A. & Blaskovich, M. A. T. (2018). Fluorescent Antibiotics: New Research Tools to Fight Antibiotic Resistance. Trends Biotechnol 36, 523–536.

[102] Jiang, M., Li, H., Johnson, A., Karasawa, T., Zhang, Y., Meier, W. B., Taghizadeh, F., Kachelmeier, A. & Steyger, P. S. (2019). Inflammation up-regulates cochlear expression of TRPV1 to potentiate drug-induced hearing loss. Sci Adv 5, eaaw1836.

[103] Ugwu, M. C., Pelis, R., Esimone, C. O. & Agu, R. U. (2017). Fluorescent organic cations for human OCT2 transporters screening: uptake in CHO cells stably expressing hOCT2. ADMET & DMPK 5, 135–145.

[104] Yasujima, T., Ohta, K., Inoue, K. & Yuasa, H. (2011). Characterization of human OCT1-mediated transport of DAPI as a fluorescent probe substrate. J Pharm Sci 100, 4006–12.

[105] Chedik, L., Bruyere, A. & Fardel, O. (2018). Interactions of organophosphorus pesticides with solute carrier (SLC) drug transporters. Xenobiotica, 1–12.

[106] Liu, X., Williams, J. B., Sumpter, B. R. & Bevensee, M. O. (2007). Inhibition of the Na/bicarbonate cotransporter NBCe1-A by diBAC oxonol dyes relative to niflumic acid and a stilbene. J Membr Biol 215, 195–204.

[107] Zou, L., Stecula, A., Gupta, A., Prasad, B., Chien, H. C., Yee, S. W., Wang, L., Unadkat, J. D., Stahl, S. H., Fenner, K. S. & Giacomini, K. M. (2018). Molecular Mechanisms for Species Differences in Organic Anion Transporter 1, OAT1: Implications for Renal Drug Toxicity. Mol Pharmacol 94, 689–699.

[108] Izumi, S., Nozaki, Y., Komori, T., Takenaka, O., Maeda, K., Kusuhara, H. & Sugiyama, Y. (2016). Investigation of Fluorescein Derivatives as Substrates of Organic Anion Transporting Polypeptide (OATP) 1B1 To Develop Sensitive Fluorescence-Based OATP1B1 Inhibition Assays. Mol Pharm 13, 438–48.

[109] Schwartz, J. W., Blakely, R. D. & DeFelice, L. J. (2003). Binding and transport in norepinephrine transporters. Real-time, spatially resolved analysis in single cells using a fluorescent substrate. J Biol Chem 278, 9768–77.

[110] Inyushin, M. U., Arencibia-Albite, F., de la Cruz, A., Vazquez-Torres, R., Colon, K., Sanabria, P. & Jimenez-Rivera, C. A. (2013). New method to visualize neurons with DAT in slices of rat VTA using fluorescent substrate for DAT, ASP+. J Neurosci Neuroeng 2, 98–103.

[111] Schwartz, J. W., Novarino, G., Piston, D. W. & DeFelice, L. J. (2005). Substrate binding stoichiometry and kinetics of the norepinephrine transporter. J Biol Chem 280, 19177–84.

[112] Haunsø, A. & Buchanan, D. (2007). Pharmacological characterization of a fluorescent uptake assay for the noradrenaline transporter. J Biomol Screen 12, 378–84.

[113] Oz, M., Libby, T., Kivell, B., Jaligam, V., Ramamoorthy, S. & Shippenberg, T. S. (2010). Real-time, spatially resolved analysis of serotonin transporter activity and regulation using the fluorescent substrate, ASP+. J Neurochem 114, 1019–29.

[114] Zwartsen, A., Verboven, A. H. A., van Kleef, R., Wijnolts, F. M. J., Westerink, R. H. S. & Hondebrink, L. (2017). Measuring inhibition of monoamine reuptake transporters by new psychoactive substances (NPS) in real-time using a high-throughput, fluorescence-based assay. Toxicol In Vitro 45, 60–71.

[115] Kido, Y., Matsson, P. & Giacomini, K. M. (2011). Profiling of a prescription drug library for potential renal drug-drug interactions mediated by the organic cation transporter 2. J Med Chem 54, 4548–58.

[116] Salomon, J. J., Endter, S., Tachon, G., Falson, F., Buckley, S. T. & Ehrhardt, C. (2012). Transport of the fluorescent organic cation 4-(4-(dimethylamino)styryl)-N-methylpyridinium iodide (ASP+) in human respiratory epithelial cells. Eur J Pharm Biopharm 81, 351–9.

[117] Rytting, E., Bryan, J., Southard, M. & Audus, K. L. (2007). Low-affinity uptake of the fluorescent organic cation 4-(4-(dimethylamino)styryl)-N-methylpyridinium iodide (4-Di-1-ASP) in BeWo cells. Biochem Pharmacol 73, 891–900.

[118] Lee, W. K., Reichold, M., Edemir, B., Ciarimboli, G., Warth, R., Koepsell, H. & Thévenod, F. (2009). Organic cation transporters OCT1, 2, and 3 mediate high-affinity transport of the mutagenic vital dye ethidium in the kidney proximal tubule. Am J Physiol Renal Physiol 296, F1504–13.

[119] Dunn, M., Henke, A., Clark, S., Kovalyova, Y., Kempadoo, K. A., Karpowicz, R. J., Jr., Kandel, E. R., Sulzer, D. & Sames, D. (2018). Designing a norepinephrine optical tracer for imaging individual noradrenergic synapses and their activity in vivo. Nat Commun 9, 2838.

[120] Masereeuw, R., Moons, M. M., Toomey, B. H., Russel, F. G. M. & Miller, D. S. (1999). Active lucifer yellow secretion in renal proximal tubule: evidence for organic anion transport system crossover. J Pharmacol Exp Ther 289, 1104–11.

[121] de Gier, R. P. E., Feitz, W. F. J., Masereeuw, R., Wouterse, A. C., Smits, D. & Russel, F. G. M. (2003). Anionic and cationic drug secretion in the isolated perfused rat kidney after neonatal surgical induction of ureteric obstruction. BJU Int 92, 452–8.

[122] Jouan, E., Le Vee, M., Denizot, C., Da Violante, G. & Fardel, O. (2014). The mitochondrial fluorescent dye rhodamine 123 is a high-affinity substrate for organic cation transporters (OCTs) 1 and 2. Fundam Clin Pharmacol 28, 65–77.

[123] Wilson, J. N., Brown, A. S., Babinchak, W. M., Ridge, C. D. & Walls, J. D. (2012). Fluorescent stilbazolium dyes as probes of the norepinephrine transporter: structural insights into substrate binding. Org Biomol Chem 10, 8710–9.

[124] Patik, I., Székely, V., Német, O., Szepesi, À., Kucsma, N., Várady, G., Szakècs, G., Bakos, È. & Özvegy-Laczka, C. (2018). Identification of novel cell-impermeant fluorescent substrates for testing the function and drug interaction of Organic Anion-Transporting Polypeptides, OATP1B1/1B3 and 2B1. Sci Rep 8, 2630.

[125] Lajiness, M. S., Maggiora, G. M. & Shanmugasundaram, V. (2004). Assessment of the consistency of medicinal chemists in reviewing sets of compounds. J Med Chem 47, 4891–6.

[126] Kell, D. B. (2013). Finding novel pharmaceuticals in the systems biology era using multiple effective drug targets, phenotypic screening, and knowledge of transporters: where drug discovery went wrong and how to fix it. FEBS J 280, 5957–5980.

[127] Kell, D. B., Wright Muelas, M., O’Hagan, S. & Day, P. J. (2018). The role of drug transporters in phenotypic screening. Drug Target Review 4, 16–19.

[128] Prior, M., Chiruta, C., Currais, A., Goldberg, J., Ramsey, J., Dargusch, R., Maher, P. A. & Schubert, D. (2014). Back to the future with phenotypic screening. ACS Chem Neurosci 5, 503–13.

[129] Swinney, D. C. (2014). Opportunities for phenotypic screening in drug discovery. Drug Disc World 15, 33–42.

[130] Yamaguchi, Y., Matsubara, Y., Ochi, T., Wakamiya, T. & Yoshida, Z. (2008). How the π conjugation length affects the fluorescence emission efficiency. J Am Chem Soc 130, 13867–9.

[131] Lavis, L. D. & Raines, R. T. (2014). Bright building blocks for chemical biology. ACS Chem Biol 9, 855–66.

[132] Zambianchi, M., Di Maria, F., Cazzato, A., Gigli, G., Piacenza, M., Della Sala, F. & Barbarella, G. (2009). Microwave-assisted synthesis of thiophene fluorophores, labeling and multilabeling of monoclonal antibodies, and long lasting staining of fixed cells. J Am Chem Soc 131, 10892–900.

[133] Di Maria, F., Palamà, I. E., Baroncini, M., Barbieri, A., Bongini, A., Bizzarri, R., Gigli, G. & Barbarella, G. (2014). Live cell cytoplasm staining and selective labeling of intracellular proteins by non-toxic cell-permeant thiophene fluorophores. Org Biomol Chem 12, 1603–10.

